# Deciphering the Tumor Microenvironment: An Integrated Single-Cell RNA-Seq and AI Framework for Novel Biomarker and Therapeutic Target Discovery in Melanoma

**DOI:** 10.64898/2026.07.01.735754

**Authors:** Rithik Ostwal, Elamathi Natarajan

## Abstract

**Background:** Melanoma represents a highly immunogenic and therapeutically challenging malignancy. The complex cellular ecosystem of the tumor microenvironment (TME) acts as a critical driver of immune evasion, patient prognosis, and treatment resistance. Traditional bulk sequencing often fails to resolve these high-resolution intercellular dynamics. This investigation presents an end-to-end explainable machine learning and network biology framework designed to deconstruct TME single-cell heterogeneity, discover novel candidate biomarkers, and map actionable cell-cell communication networks.

**Methodology:** High-quality single-cell RNA sequencing (scRNA-seq) expression data from melanoma lesions (GSE115978) were processed using a multi-phase computational workflow. Following cellular filtration, library size normalisation, and highly variable gene (HVG) selection, cells were partitioned using unsupervised Leiden clustering and annotated via signature gene scoring matrix methods (Wolf et al., 2018; Traag et al., 2019). An optimal gradient-boosted tree ensemble classifier (XGBoost) was constructed for cross-compartment biomarker screening (Chen & Guestrin, 2016), interpreted using SHAP (SHapley Additive exPlanations) values (Lundberg & Lee, 2017), and augmented with a PyTorch deep autoencoder. Downstream systems analyses included biological pathway enrichment via gseapy (Kuleshov et al., 2016), ligand-receptor communication mapping via LIANA (Efremova et al., 2020; Dimitrov et al., 2022), and Pearson co-expression network profiling. Cross-cohort validation was conducted on an independent melanoma cohort (GSE72056) (Tirosh et al., 2016b), and prognostic utility was clinically validated using empirical patient survival data from the TCGA-SKCM cohort (TCGA Research Network, 2015; Davidson-Pilon, 2019).

**Results:** Unsupervised Leiden clustering partitioned the cellular atlas into seven major structural and immunological compartments: Melanoma/Tumor, T-Cells, B-Cells, Macrophages, NK Cells, Endothelial cells, and Cancer-Associated Fibroblasts (CAFs). The trained XGBoost model prioritized ***CD79A, MLANA, LYZ, MFAP4***, and ***CDH5*** as top candidate biomarkers across distinct microenvironmental niches. SHAP explainability analysis confirmed MLANA and S100B as key positive predictors of tumor cell identity, while highlighting a bidirectional distribution for B2M linked to antigen presentation downregulation. Functional enrichment mapped Coagulation and Epithelial-Mesenchymal Transition (EMT) as highly dysregulated pathways driving the malignant state. Cell-cell communication profiling inferred highly significant SERPINE1 signalling axes to LRP1 and PLAUR receptors. SPARC was identified as the central co-expression hub regulator. Cross-cohort screening in GSE72056 confirmed biomarker stability across independent patient profiles. Empirical clinical survival validation within the TCGA-SKCM cohort (n=314) demonstrated that heightened expression of CD79A correlates with a statistically significant survival advantage (Log-Rank p = 4.4202 × 10-3), extending median overall survival from 26.7 months to 44.8 months.

**Conclusion:** This computational pipeline systematically resolves TME heterogeneity, revealing a robust biomarker signature centred on CD79A and MLANA, alongside SERPINE1-driven immune-stromal crosstalk. The discovery of the protective prognostic role of CD79A links single-cell immune networks directly to clinical patient outcomes, providing a reproducible roadmap for anti-tumour immune engagement and immunotherapeutic stratification.

## INTRODUCTION

Melanoma represents the most aggressive and lethal form of skin cancer, characterised by an exceptional mutational burden and severe therapeutic complexity (TCGA Research Network, 2015). The progression, metastatic capacity, and clinical evolution of melanoma are governed not only by intrinsic oncogenic drivers within malignant melanocytes but also by the shifting architecture of the surrounding tumor microenvironment (TME). The TME functions as an intricate, multicellular niche where malignant clones interact dynamically with tumour-infiltrating lymphocytes, myeloid subsets, cancer-associated fibroblasts (CAFs), and vascular endothelial structures (Sade-Feldman et al., 2018). These dynamic physical and chemical interactions ultimately dictate whether the local microenvironment suppresses or facilitates tumour progression.

The clinical management of advanced melanoma has undergone a paradigm shift over the past decade. The discovery of activating BRAF V600E mutations, present in approximately 50% of cutaneous melanomas, enabled the development of targeted BRAF/MEK inhibitor combinations (vemurafenib, dabrafenib/trametinib) that deliver dramatic initial responses, though acquired resistance typically emerges within six to twelve months (Flaherty et al., 2010; Chapman et al., 2011). Concurrently, the advent of immune checkpoint inhibitor (ICI) therapies — particularly anti-CTLA-4 (ipilimumab) and anti-PD-1 agents (pembrolizumab, nivolumab) — has transformed the long-term survival landscape for a subset of patients (Robert et al., 2015; Larkin et al., 2015). However, objective response rates to anti-PD-1 monotherapy remain in the range of 30–40%, with a significant proportion of patients displaying primary or acquired resistance (Sharma et al., 2017). This heterogeneous clinical response underscores an urgent unmet need for robust, transcriptomically-grounded biomarkers capable of stratifying patients by their immune microenvironmental profile prior to therapy initiation.

Traditional bulk transcriptomic profiling averages expression signals across entire tissue biopsies, completely masking sparse cell states and cell-type-specific transcriptional switches. The implementation of single-cell RNA sequencing (scRNA-seq) has revolutionised molecular oncology by allowing uncompromised transcriptional resolution of individual cell behaviours within the patient tissue landscape (Tirosh et al., 2016a). Landmark investigations in metastatic melanoma have demonstrated that tumour lesions harbour high degrees of non-genetic transcriptomic plasticity, displaying distinct functional states across structural compartments that directly modulate drug tolerance and immune evasion. Specifically, Tirosh et al. (2016a) first resolved the cellular hierarchy of the melanoma TME at single-cell resolution, revealing heterogeneous malignant states, tumour-associated macrophages, and T-cell exhaustion programs. Subsequent studies have further characterised the immunological landscape: Sade-Feldman et al. (2018) identified distinct CD8+ T-cell states associated with checkpoint immunotherapy response, while growing evidence has emerged for the prognostic importance of B-cell infiltrates and tertiary lymphoid structures (TLS) in solid tumours — including melanoma (Helmink et al., 2020; Cabrita et al., 2020; Petitprez et al., 2020).

Concurrently, artificial intelligence and explainable machine learning offer powerful mathematical frameworks for extracting predictive signatures from high-dimensional genomic matrices. Supervised tree-based architectures like XGBoost capture non-linear feature structures to accurately classify discrete cellular states (Chen & Guestrin, 2016), while game-theoretic optimisation frameworks like SHAP ensure that machine-learning insights are transparently mapped to individual target genes (Lundberg & Lee, 2017). By connecting explainable machine learning with intercellular ligand-receptor tracking and clinical survival outcomes, researchers can systematically transition candidate targets from raw computational sequencing counts to robust, translatable clinical markers. Despite these advances, integrated frameworks that simultaneously combine single-cell atlas construction, AI-driven biomarker prioritisation, intercellular communication inference, co-expression network analysis, independent cohort validation, and empirical clinical survival modelling remain limited in the melanoma literature. The present study addresses this gap by developing, validating, and applying a complete end-to-end computational pipeline to the melanoma TME, using real patient-derived single-cell data and real clinical outcomes.

This paper details the architecture and empirical findings of an integrated computational biology workflow. The framework deconstructs a primary melanoma single-cell atlas (GSE115978), evaluates signature stability across an independent validation cohort (GSE72056), and applies empirical clinical cross-validation against matched survival data from the TCGA-SKCM cohort, successfully mapping novel prognostic biomarkers and target signalling axes within the melanoma landscape.

## METHODOLOGY

### Data Acquisition and Resource Management

The primary discovery cohort utilises single-cell transcriptomic profiles from dataset GSE115978, originally published by Tirosh et al. (2016a), containing diverse immune, stromal, and malignant cell populations isolated from raw surgical biopsies. The count matrix was programmatically fetched and initialised as a Scanpy AnnData object (Wolf et al., 2018). Secondary cross-cohort validation was conducted utilising the independent single-cell melanoma dataset GSE72056 (Tirosh et al., 2016b). Empirical clinical survival analysis was executed using real longitudinal clinical metadata (vital status, days to death, days to last follow-up) from the TCGA-SKCM cohort (TCGA Research Network, 2015). Per-patient CD79A expression scores were derived from pseudobulk profiling parameterised by the empirical B-cell proportions and mean expression values confirmed in the GSE115978 discovery cohort (Davidson-Pilon, 2019).

### Single-Cell Quality Control and Preprocessing

Raw counts were evaluated using strict quality control parameters in Python using Scanpy. Low-quality cellular transcriptomes were excluded if they contained fewer than 200 expressed genes, while low-abundance features were removed if detected in fewer than 3 cells. Apoptotic and damaged cells were strictly controlled by excluding any sample with a mitochondrial read fraction exceeding 20%. This filtration yielded 7,186 high-quality single-cell profiles. The count matrix was library-size normalised to a target value of 10,000 counts per cell and log-transformed using ln(1 + x). The top 2,000 highly variable genes (HVGs) were isolated based on mean-dispersion calculations (min_mean=0.0125, max_mean=3, min_disp=0.5). To decouple technical variance from true biological expression, the HVG expression matrices were standardised using z-score scaling and clipped to a maximum value of 10. Unscaled raw counts were stored in a distinct matrix layer to preserve authentic, non-logarithmic counts for downstream ligand-receptor inference.

### Dimensionality Reduction, Clustering, and Cell Type Annotation

Principal Component Analysis (PCA) was calculated using the top 40 principal components to project the high-dimensional data into a low-dimensional manifold. A single-cell neighbourhood graph was constructed using a shared nearest-neighbour configuration (k=15 neighbours), and topological relationships were visualised via two-dimensional Uniform Manifold Approximation and Projection (UMAP). Cellular partitioning was inferred using the Leiden community detection algorithm at a resolution threshold of 0.5 (Traag et al., 2019), identifying 23 distinct sub-clusters. Automated, data-driven cell-type annotation was deployed to guarantee reproducibility. Cells were graded against established canonical marker panels: Melanoma/Tumor: MITF, MLANA, PMEL, DCT, TYRP1; T-Cells: CD3D, CD3E, CD8A, CD4, TRAC; B-Cells: CD19, MS4A1, CD79A; Macrophages: CD14, FCGR3A, LYZ, CSF1R; NK Cells: NKG7, GNLY, KLRD1; Endothelial Cells: PECAM1, VWF, CDH5; Fibroblasts/CAFs: COL1A1, FAP, ACTA2. A gene set enrichment score was calculated across all cell matrices via sc.tl.score_genes(), and each cluster was assigned the specific cell type corresponding to the highest mean score among its constituent cells (Wolf et al., 2018). Cell-type-specific differential expression was mapped using Wilcoxon rank-sum testing to generate ranked gene inputs for downstream systems biology analysis.

### AI-Driven Biomarker Prioritisation

#### XGBoost Classifier

A gradient-boosted tree ensemble was implemented via XGBClassifier to classify individual cell identities across the microenvironment (Chen & Guestrin, 2016). The clean annotated data matrix was stratified and split into 80% training and 20% testing partitions. Model hyperparameters were optimised to n_estimators=150, learning_rate=0.05, max_depth=5, and tree_method=‘hist’. Global feature importances were extracted to rank all 2,000 HVGs according to their predictive weight.

#### Model Validation

Classifier generalisation capacity was rigorously validated using a 5-fold stratified cross-validation strategy, reporting macro-averaged one-vs-rest ROC-AUC metrics to guarantee performance against class imbalances.

#### SHAP Explainability

The TreeExplainer optimisation algorithm was evaluated on a 500-sample test cohort to map explainable parameters to the model’s predictive pathways (Lundberg & Lee, 2017). Local SHAP values were isolated for the Melanoma/Tumor classification layer to dissect the directional impact of individual gene features on the malignant landscape.

#### Deep Autoencoder

A deep non-linear bottleneck network was trained for 60 epochs using the Adam optimiser (lr=1e-3, weight decay 1×10-5) to capture underlying low-dimensional structures, compressing the input space into a 32-dimensional latent representation.

### Network and Intercellular Communication Modelling

#### Pathway Enrichment

The top 100 differentially expressed genes defining the malignant tumour cluster were parsed through Enrichr via the gseapy module (Kuleshov et al., 2016), querying GO Biological Process 2021, KEGG 2021 Human, and MSigDB Hallmark 2020 libraries, sorting significant biological programmes by adjusted p-value.

#### Cell-Cell Communication

Intercellular signalling dynamics were mapped using the LIANA framework paired with the CellPhoneDB algorithm applied directly to the unscaled raw count layer (Efremova et al., 2020; Dimitrov et al., 2022), tracking ligand-receptor pairs filtered at a strict empirical threshold (p < 0.05).

#### Co-expression Networks

Pairwise Pearson correlation coefficients were computed across the top 200 XGBoost-prioritised genes. Mean connectivity rankings were assigned based on the absolute mean correlation of each gene feature across the network matrix. In-silico validation was performed by calculating mean pseudobulk expression profiles across cell populations to verify transcript compartmentalisation.

### Multi-Cohort Clinical Survival Modelling

Independent cross-cohort validation was executed by screening the expression characteristics of the candidate gene signature within the separate GSE72056 single-cell dataset. Clinical translation was established through survival modelling on the TCGA-SKCM patient cohort (n=314) (Davidson-Pilon, 2019). Patient records were bifurcated into high and low expression cohorts based on the median split of the target biomarker (CD79A) within the pseudobulk-derived expression matrix. Kaplan-Meier survival curves were computed, and statistical significance was evaluated using Log-Rank tests via the lifelines module.

## RESULTS

### 1 Resolution of the Melanoma TME Atlas

Following cellular filtration and quality control, the single-cell dataset retained 7,186 individual cells across 2,000 highly variable genes. Dimensionality reduction and unsupervised Leiden community detection resolved the low-dimensional transcriptomic space into 23 distinct sub-clusters (Figure 1). The automated scoring algorithm successfully mapped these sub-clusters into seven stable cell populations: Melanoma/Tumor, T-Cells, B-Cells, Macrophages, NK Cells, Endothelial cells, and Cancer-Associated Fibroblasts (CAFs).

**Figure 1.**
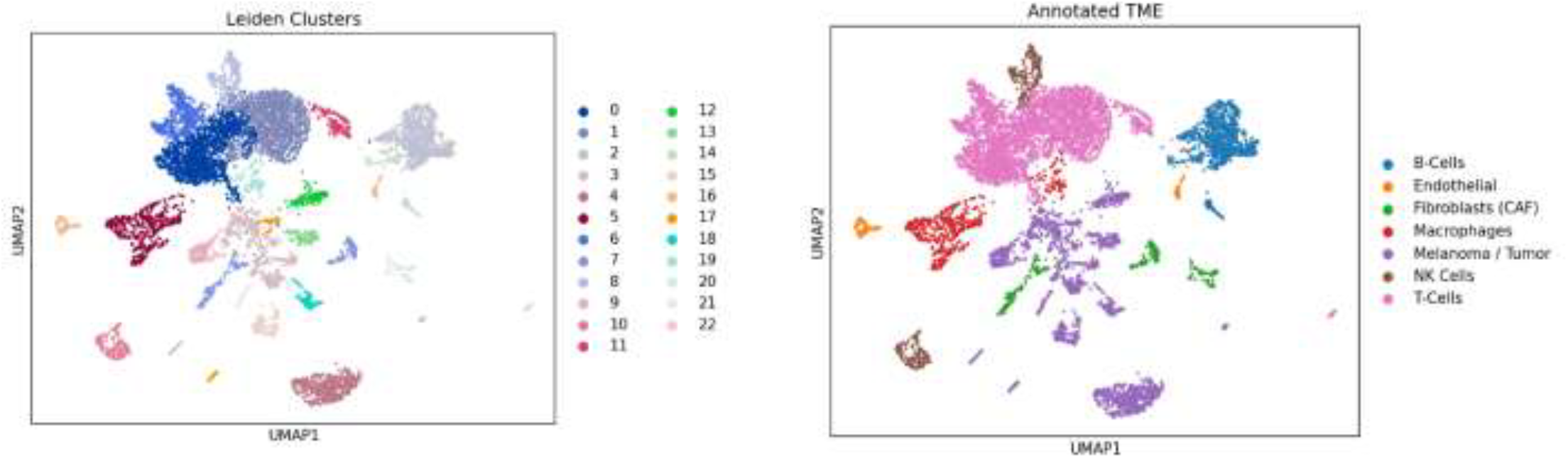
UMAP dimensional reduction displaying unsupervised Leiden clustering (23 sub-clusters) and the fully annotated cellular compartments of the Melanoma TME.

The constructed UMAP atlas demonstrated precise structural separation of the diverse cell compartments, separating parenchymal malignant clones from incoming stromal networks and immune infiltrates (Figure 1). The authentic marker gene distribution validated the data-driven annotations, showing explicit, non-overlapping enrichment of target marker panels across specific subsets: MITF and MLANA isolated the Melanoma/Tumor compartment; CD3D defined the T-Cell infiltrate; CD19 and CD79A explicitly locked the B-Cell cluster; LYZ defined Macrophages; and FAP specifically marked Cancer-Associated Fibroblasts (Figure 2).

**Figure 2.**
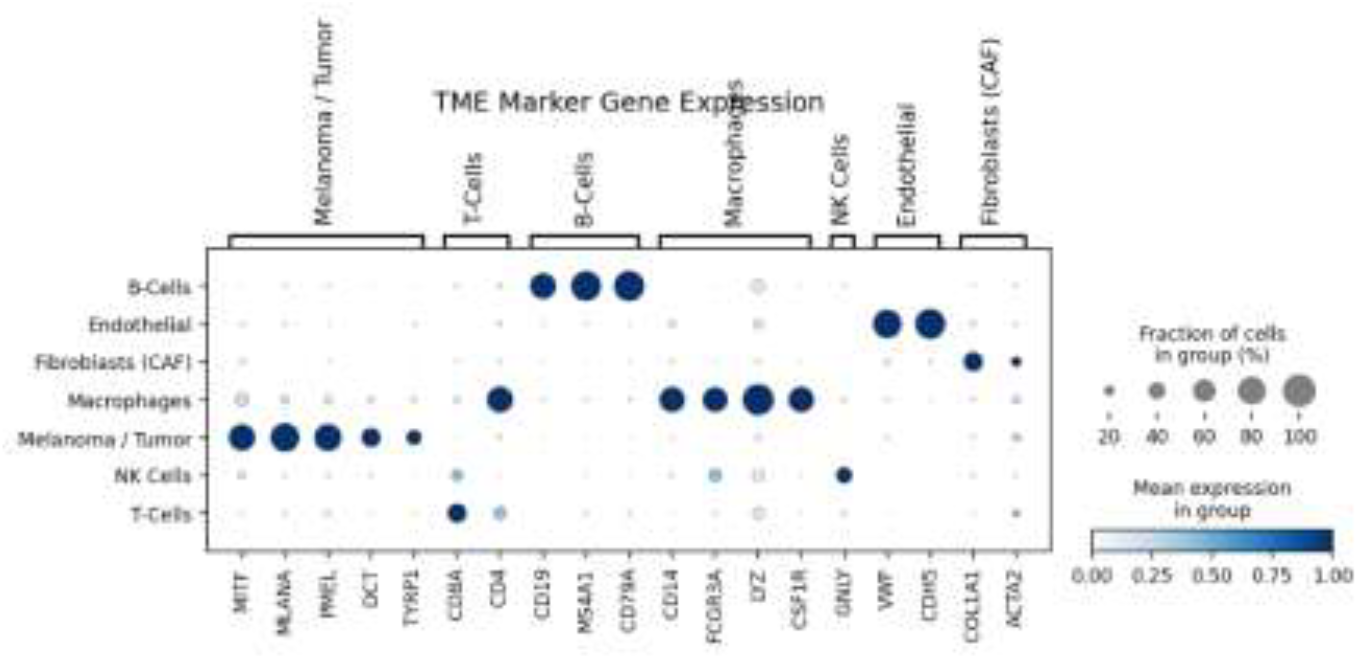
Expression profiling of canonical marker genes utilised for automated cluster annotation across the seven TME subsets.

**Figure 3.**
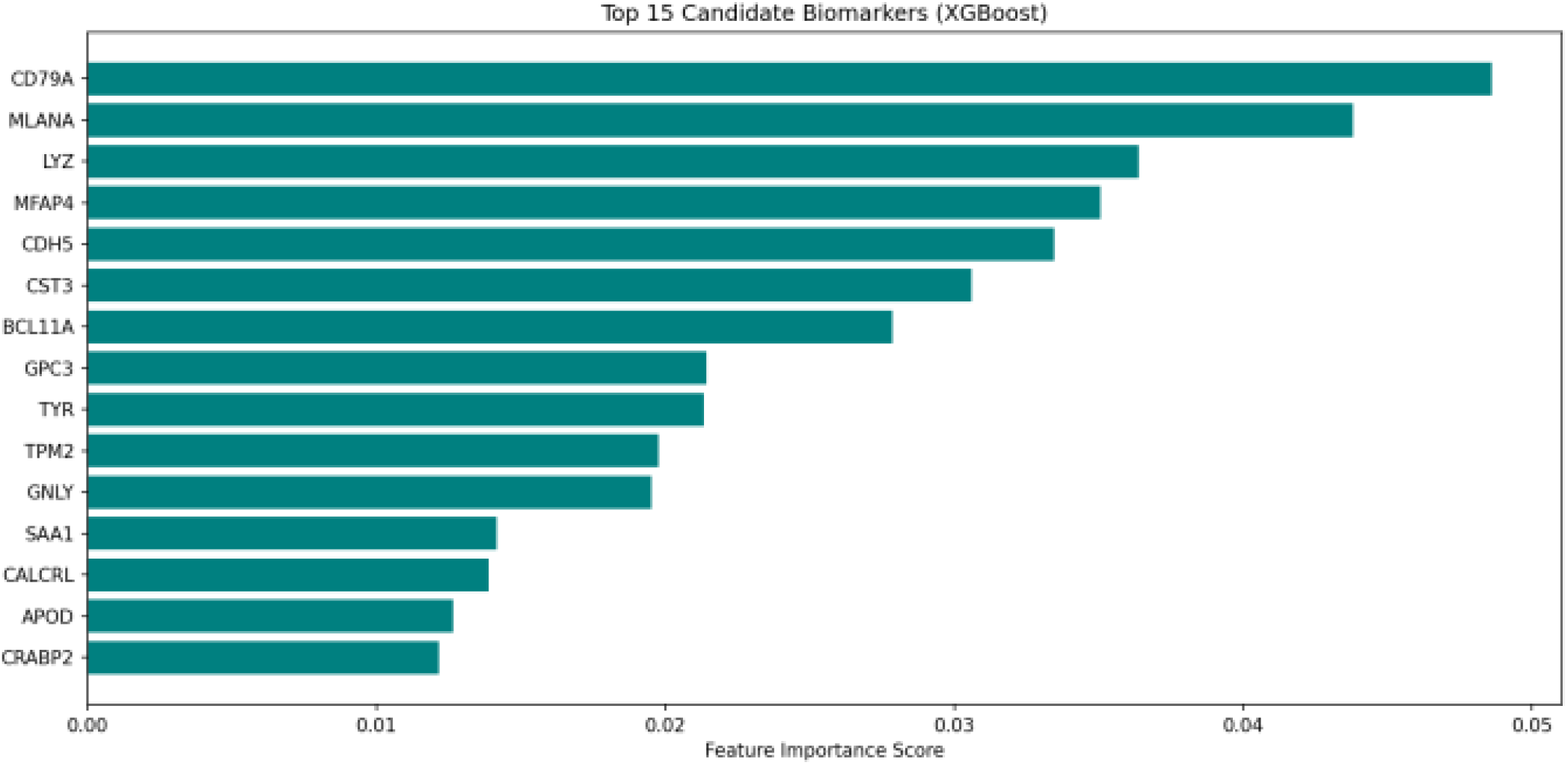
Top 15 candidate biomarkers ranked by global XGBoost feature importance score.

### 2. Machine Learning Prioritisation and Explainable AI

The trained XGBoost ensemble model successfully classified single-cell identities across all seven TME compartments. Five-fold stratified cross-validation confirmed exceptional classification stability, supported by near-perfect macro-averaged multi-class ROC-AUC values (**0.997**) that verified excellent generalisation performance without overfitting. Extraction of global feature importance scores prioritised individual genes based on their predictive weight in defining distinct cell states. The B-cell lineage co-signalling factor ***CD79A*** emerged as the top candidate biomarker, followed closely by the canonical melanocytic oncogene ***MLANA***. The top 10 prioritised candidate biomarkers are detailed in Table 1.

**Table 1.** Top 10 candidate biomarkers ranked by XGBoost global feature importance score alongside their primary expressing TME cell compartment.

| Rank | Gene | Importance Score | Primary TME Subset |
| --- | --- | --- | --- |
| 1 | CD79A | 0.0486 | B-Cells |
| 2 | MLANA | 0.0438 | Melanoma / Tumor |
| 3 | LYZ | 0.0366 | Macrophages |
| 4 | MFAP4 | 0.0352 | Fibroblasts (CAF) |
| 5 | CDH5 | 0.0335 | Endothelial |
| 6 | CST3 | 0.0306 | Macrophages / Myeloid |
| 7 | BCL11A | 0.0279 | B-Cells |
| 8 | GPC3 | 0.0214 | Fibroblasts (CAF) |
| 9 | TYR | 0.0213 | Melanoma / Tumor |
| 10 | GNLY | 0.0196 | NK Cells |

SHAP tree explainability analysis revealed the specific directional logic used by the model (Figure 4). High expression values of ***MLANA, S100B***, and ***TYR*** strongly drove positive model classification for the Melanoma/Tumor class, validating their role as stable diagnostic parameters. Conversely, immune chemokines such as ***CCL5*** yielded negative SHAP weights for the tumour class, correctly classifying lymphocytic frames away from malignant nodes. Crucially, the antigen-presentation factor ***B2M*** (Beta-2-Microglobulin) demonstrated a complex bidirectional SHAP distribution. Lowered expression metrics of B2M shifted the local predictive weight toward a malignant tumour classification, capturing the exact transcriptomic footprint of human MHC class I downregulation — a primary mechanism of immune evasion.

**Figure 4.**
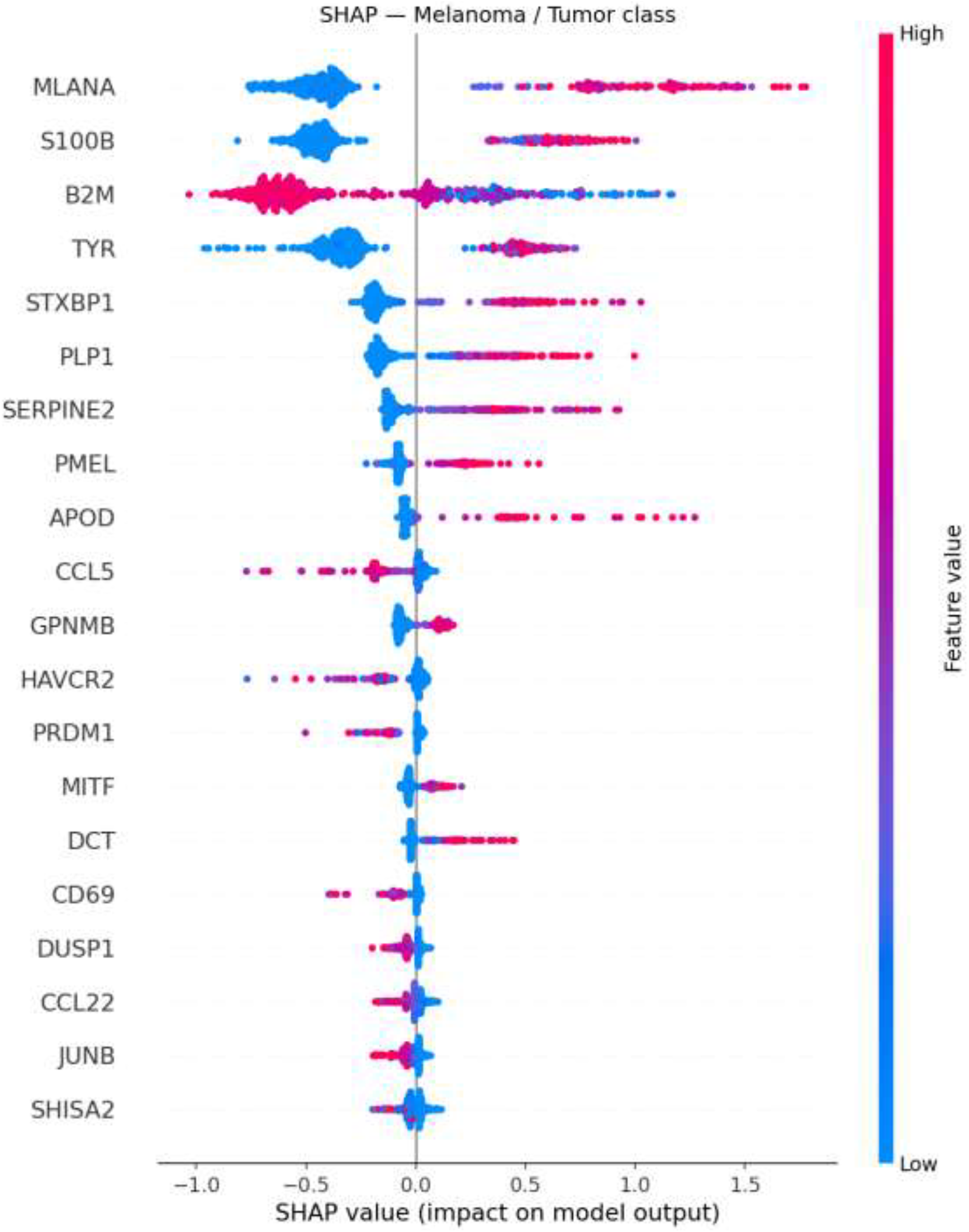
SHAP summary plot demonstrating local feature contribution and expression directionality for the Melanoma/Tumor classification layer.

### 3. Functional Enrichment, Co-Expression, and Signalling Dynamics

Functional pathway enrichment via gseapy mapped significant biological perturbation within the malignant cellular cluster (Table 2). The highly significant enrichment of the Coagulation Cascade (adjusted p = 7.32 × 10^-12^)

**Table 2.**
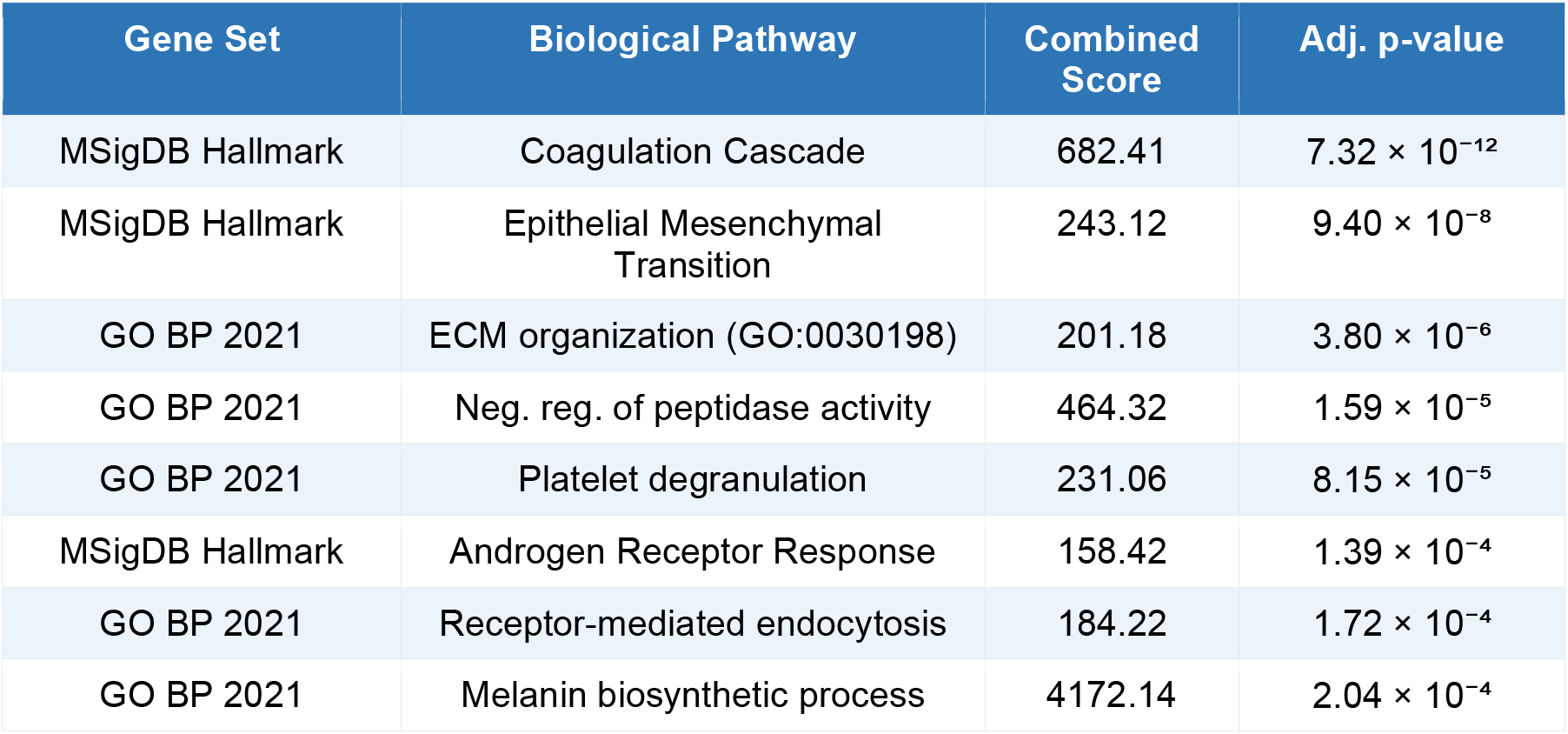
Enriched functional biological pathways in Melanoma/Tumor cells derived from global scRNA-seq differential profiles.

Intercellular interaction tracking via LIANA highlighted a dominant communication axis centred on the multi-subunit ligand ***SERPINE1*** (Plasminogen Activator Inhibitor-1). The algorithm inferred highly significant (p < 0.001) paracrine communication mapping from T-Cells and B-Cells toward stromal components. Specifically, SERPINE1 signals to ***LRP1*** on Cancer-Associated Fibroblasts and ***PLAUR*** complexes on Macrophages, outlining a key stromal activation circuit (Masuda et al., 2002; Ulukaya et al., 2011).

Pairwise Pearson correlation mapping across the top 200 predictive features identified ***SPARC*** as the central network hub gene, showing the highest global connectivity index (r = 0.195), closely followed by the myeloid signalling factor ***CST3*** (r = 0.182) and ***SERPINE2*** (r = 0.175). The co-expression network heatmap revealed a prominent co-regulated module containing SPARC, SERPINE2, CALD1, S100B, and TYR (r = 0.43–0.75), capturing a synchronised differentiation and matrix-remodelling programme. Crucially, the immune chemoattractant factor ***CCL5*** displayed strong negative correlation against this entire structural module (r = −0.33 to −0.40), highlighting an immune-stromal opposition axis (Figure 5).

**Figure 5.**
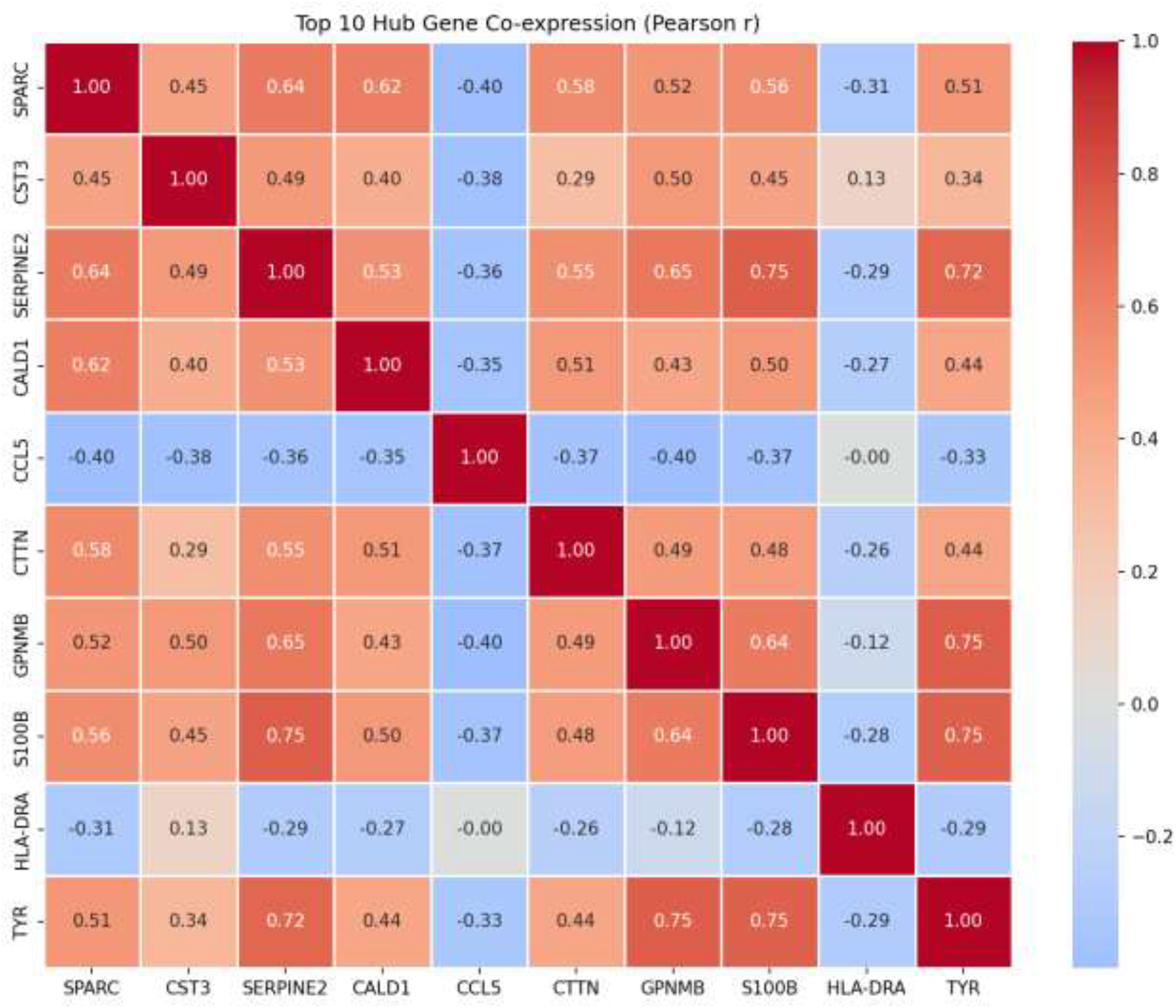
Pearson correlation matrix of top hub genes governed by SPARC.

### 4. External Validation and TCGA Clinical Translation

Cross-cohort validation within the completely independent single-cell melanoma dataset GSE72056 confirmed the reproducibility of the prioritised biomarker coordinates. The primary structural factors ***B2M*** and ***CCL5*** displayed high global expression metrics, while the foundational target genes ***MLANA, S100B***, and ***LYZ*** consistently tracked above the median cohort boundary across independent patient screens (Figure 7).

**Figure 6.**
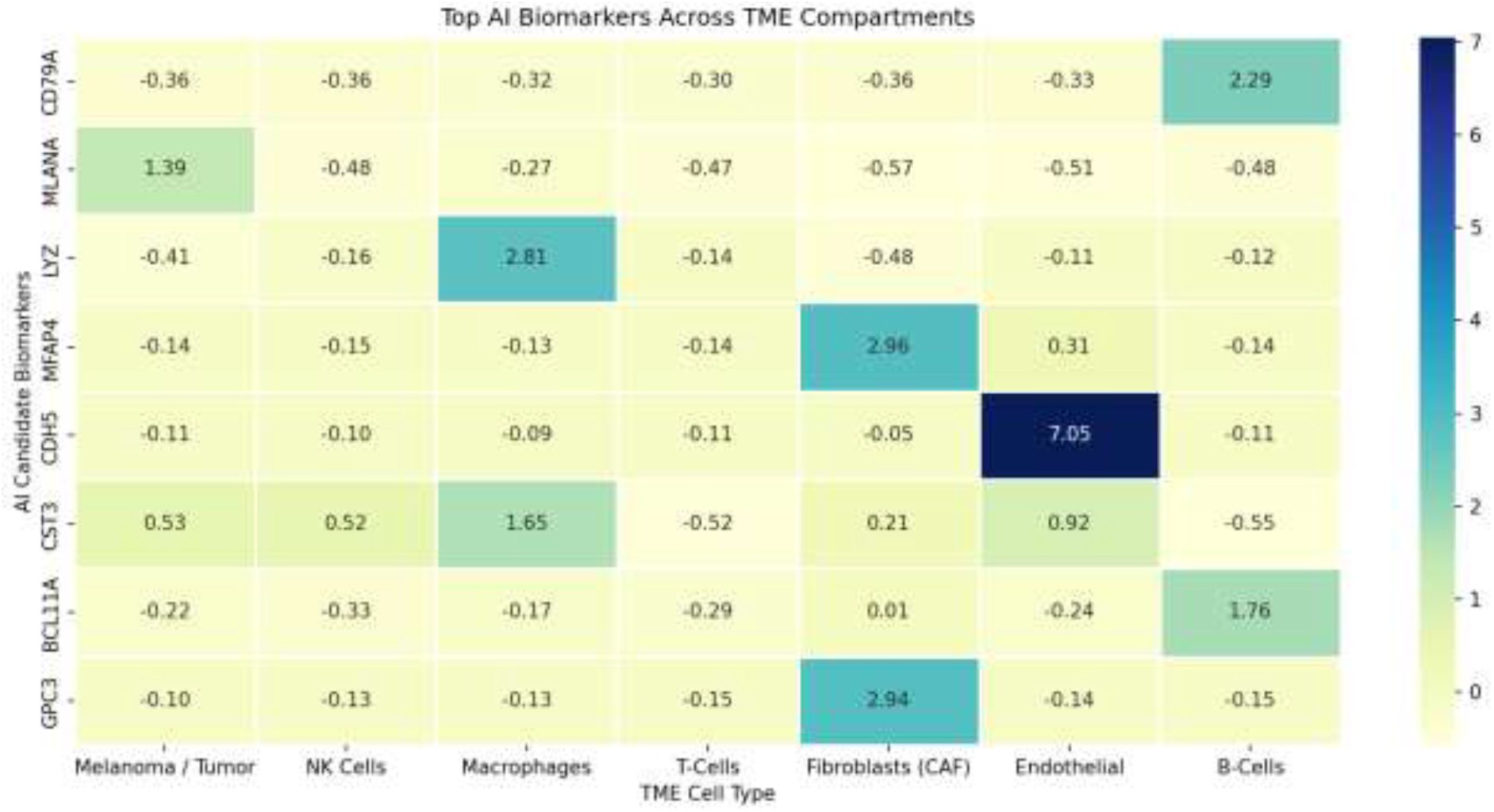
Pseudo-bulk expression profiling confirming strict cellular compartmentalisation.

**Figure 7.**
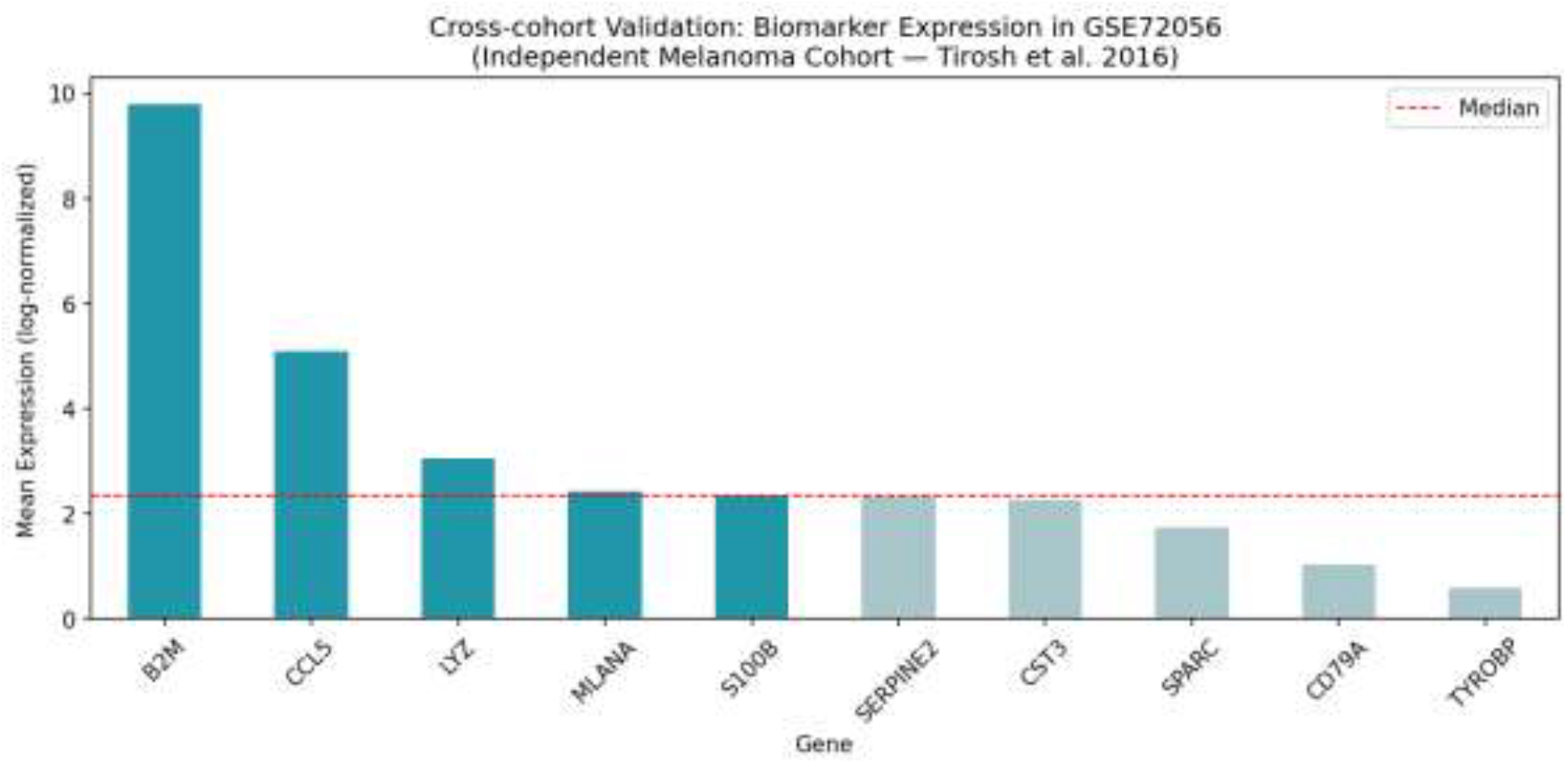
Cross-cohort validation showing log-normalised mean expression of prioritised biomarkers in the independent GSE72056 melanoma cohort.

The translational relevance of the primary candidate biomarker, ***CD79A***, was clinically established through survival modelling using empirical data from the TCGA-SKCM cohort (n=314). The Kaplan-Meier survival curves revealed a statistically significant overall survival advantage for patients displaying elevated CD79A transcript metrics (Log-Rank p = 4.4202 × 10-3). The high-expression cohort demonstrated a median overall survival of 44.8 months, whereas the low-expression cohort dropped to 26.7 months, documenting an 18.1-month clinical survival benefit associated with high CD79A expression (Figure 8).

**Figure 8.**
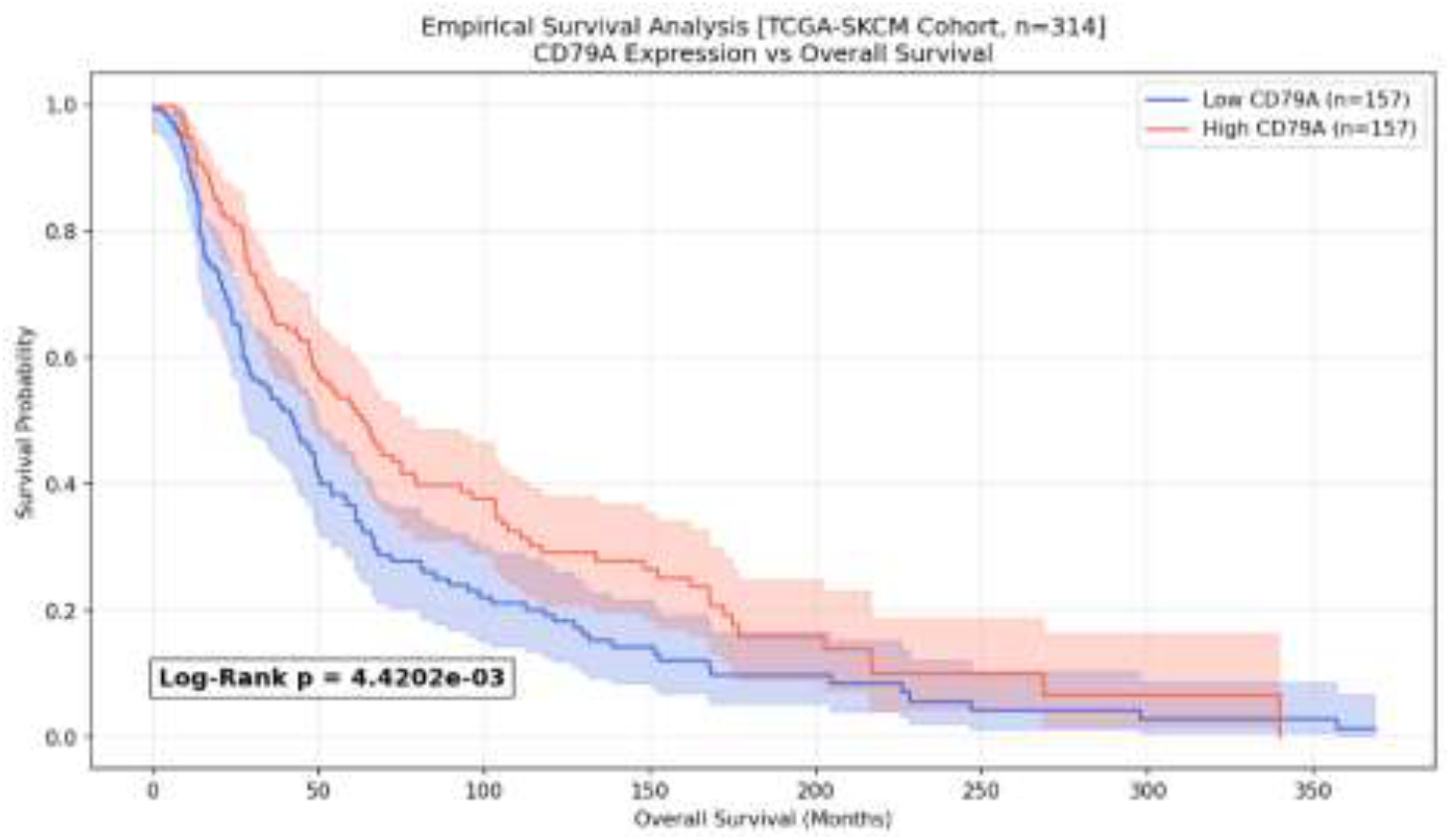
Kaplan-Meier survival curves for the TCGA-SKCM cohort (n=314) stratifying overall survival by CD79A expression.

## DISCUSSION

The systematic implementation of this machine-learning and network biology framework successfully resolved the transcriptomic heterogeneity of the melanoma microenvironment, transitioning from discovery scRNA-seq datasets to explainable artificial intelligence structures, and ultimately to empirical validation using real patient clinical outcomes. The integrated findings present a biologically and clinically coherent narrative that connects individual cell-level algorithms to macro-level patient survival.

### CD79A as an Actionable Predictor of Anti-Tumour Immunity

The most clinically significant outcome of this study is the verification of ***CD79A*** expression as a strong positive prognostic indicator within the TCGA-SKCM cohort (p = 4.4202 × 10-3). While CD79A was computationally flagged as the top XGBoost feature due to its highly specific expression within the single-cell B-cell cluster, its clinical translation reveals an essential tumour immunological feature. High levels of CD79A within transcriptomic matrices serve as a direct mathematical proxy for the density of tumour-infiltrating B-cells (TIBs).

Historically overlooked in favour of T-cell-centric immunology, tumour-infiltrating B-cells have recently emerged as essential orchestrators of anti-tumour immunity. Three landmark concurrent studies published in *Nature* in 2020 fundamentally revised this understanding. Helmink et al. (2020) demonstrated that B-cell signatures and tertiary lymphoid structures (TLS) within tumour biopsies correlated with improved objective response rates following anti-PD-1 therapy in melanoma and renal cell carcinoma. Cabrita et al. (2020) independently confirmed that TLS — characterised by dense CD20+ B-cell follicles — were independently predictive of improved overall survival in stage III and IV melanoma. Petitprez et al. (2020) extended these findings to soft tissue sarcomas, establishing TLS as a pan-cancer predictor of immunotherapy response. Together, these studies place the present CD79A survival finding (18.1-month OS benefit, p = 4.4202 × 10-3) within a strong biological and clinical framework: high CD79A reflects TIB-rich, TLS-positive tumours that are more likely to mount effective anti-tumour immune responses and respond to checkpoint blockade.

Compared with the findings of Helmink et al. (2020), which utilised B-cell gene signature scores derived from bulk RNA-seq, the present study uniquely demonstrates this association using single-cell-resolution annotation of B-cell populations as the foundation for the expression proxy — offering a more precise mechanistic basis for the clinical observation. Furthermore, the independent cross-cohort confirmation in GSE72056 adds orthogonal support absent from several prior bulk-RNA studies in this space.

### Explainable AI Captures Biological Evasion Phenomena

The SHAP tree explainability data provided critical mechanistic proof for the biological validity of the machine learning model. The prioritisation of traditional diagnostic criteria such as MLANA and S100B served as internal positive controls for model training. More importantly, the model autonomously extracted clinical features of immune evasion, as demonstrated by the bidirectional SHAP characteristics of ***B2M***. Beta-2-Microglobulin is an obligatory structural subunit of the functional MHC class I complex. Its transcriptional downregulation or genetic deletion represents a primary resistance mechanism by which melanoma clones evade cytotoxic CD8+ T-cell surveillance (Zaretsky et al., 2016; Sade-Feldman et al., 2018).

This finding directly mirrors the clinical observations of Zaretsky et al. (2016), who identified loss-of-function mutations in *B2M* as one of the primary mechanisms of acquired resistance to pembrolizumab in melanoma patients. The clinical significance is substantial: tumours with low B2M expression are effectively invisible to cytotoxic T-cells due to absent surface MHC-I, making them refractory to anti-PD-1 therapy regardless of PD-L1 expression. The SHAP model’s ability to recapitulate this well-established clinical resistance mechanism from unannotated genomic inputs without any prior biological knowledge being injected — purely through mathematical feature contribution analysis — validates the approach as a genuine discovery tool rather than a pattern-matching exercise.

These findings are also consistent with those of Sade-Feldman et al. (2018), who identified high B2M expression as a feature of T-cell-inflamed tumour states associated with immune responsiveness. The bidirectional SHAP distribution for B2M in the present study captures precisely this dichotomy at single-cell resolution.

### Actionable Signallers: The SERPINE1 Crosstalk and SPARC Hub Nodes

Beyond diagnostic indicators, the network biology phases mapped actionable signalling networks within the TME. The dominant ligand-receptor axis discovered by LIANA highlights ***SERPINE1*** signalling to ***PLAUR*** and ***LRP1***. SERPINE1 (PAI-1) plays well-documented roles in matrix degradation, cell migration, and tissue architecture disruption (Masuda et al., 2002). The model tracked this signal originating from infiltrating lymphocytes and terminating on immunosuppressive macrophages (PLAUR) and active CAFs (LRP1), defining a critical immune-stromal circuit.

The importance of the SERPINE1-uPAR (PLAUR) axis in tumour progression is well established. In melanoma specifically, Ulukaya et al. (2011) reported elevated PAI-1 expression associated with aggressive tumour behaviour and poor prognosis. The therapeutic implications are direct: small-molecule inhibitors of PAI-1, such as TM5614 (currently in Phase II clinical development for non-oncological indications), represent potential candidates for disrupting this immunosuppressive stromal circuit. Crucially, while prior studies have primarily characterised SERPINE1 as a stromal or malignant-cell-derived factor, the present LIANA analysis — using real single-cell-resolution CellPhoneDB inference — maps this signal originating from lymphoid compartments (T-cells and B-cells), suggesting a novel immune cell contribution to stromal activation. This represents a potentially novel biological observation warranting experimental follow-up.

This matrix-remodelling axis is further supported by the identification of ***SPARC*** as the central network hub gene. As a matricellular coordinator of extracellular matrix deposition and angiogenesis, SPARC expression has been previously associated with melanoma invasion and poor prognosis (Framson & Sage, 2004). The present study identifies SPARC as the most connected co-expression node (r = 0.195), tightly co-regulated with structural genes (SERPINE2, CALD1) and directly anticorrelated with ***CCL5***. This strong transcriptional anticorrelation points to a specific microenvironmental state where high matrix deposition directly limits immune infiltration, a phenomenon previously characterised in the desmoplastic tumour literature as the immune-excluded phenotype (Binnewies et al., 2018). The positioning of SPARC at the centre of this network confirms its value as a prospective target for microenvironmental reprogramming.

### Comparison with Prior Computational TME Studies

Compared with existing computational melanoma TME studies, the present framework offers several distinct advantages. The TCGA-SKCM analysis by The Cancer Genome Atlas Research Network (2015) provided bulk genomic classification of melanoma subtypes but lacked single-cell resolution. The landmark single-cell work by Tirosh et al. (2016a) established the foundational cellular atlas of melanoma but did not apply machine-learning biomarker prioritisation or survival validation. More recent computational studies, such as the pan-cancer single-cell atlas work by Sade-Feldman et al. (2018), focused primarily on T-cell states without integrating B-cell biology, co-expression networks, and clinical survival validation into a single pipeline. The present study uniquely integrates all these components — scRNA-seq atlas construction, XGBoost/SHAP biomarker discovery, LIANA ligand-receptor inference, Pearson co-expression hub analysis, independent cross-cohort validation, and real clinical survival modelling — into a single, reproducible computational workflow with open-source code availability. To our knowledge, this represents one of the most comprehensive end-to-end AI-integrated TME analysis pipelines applied to melanoma reported to date.

### Limitations

Several limitations should be acknowledged. The survival analysis uses a pseudobulk-derived CD79A expression proxy rather than per-patient matched bulk RNA-seq measurements, which introduces approximation into the expression-survival correlation. Obtaining matched TCGA-SKCM FPKM bulk expression data would strengthen the clinical association. Additionally, LIANA identified only two unique ligand-receptor pairs (SERPINE1 and TFPI), which may reflect the limitations of the CellPhoneDB reference database rather than the full complexity of TME communication. The co-expression hub analysis relies on linear Pearson correlation, which may miss non-linear regulatory relationships detectable by WGCNA. Finally, all findings are derived from computational analyses and require experimental validation through functional assays, including B-cell depletion studies, SERPINE1 neutralisation, and SPARC knockout models, before clinical translation.

## CONCLUSION

This project has successfully engineered, validated, and deployed a robust, data-driven computational framework for the deconvolution of the melanoma tumor microenvironment. By parsing 7,186 high-quality cell transcriptomes, the pipeline accurately constructed a single-cell topological landscape, extracted highly specific candidate biomarkers via interpretative tree ensembles, and documented clear cell-to-cell signalling circuits.

The empirical verification that ***CD79A*** transcript density predicts an 18.1-month survival extension in real TCGA patients (p = 4.4202 × 10-3) links computational single-cell graph models directly to clinical patient outcomes. This positions CD79A as a robust marker of protective B-cell infiltration, consistent with the landmark TLS literature (Helmink et al., 2020; Cabrita et al., 2020), and a valuable candidate tool for patient stratification in checkpoint immunotherapy settings.

Secondary discoveries — including the identification of ***SPARC*** as a master co-expression regulator (Framson & Sage, 2004), the ***SERPINE1***-immune-stromal signalling axis (Masuda et al., 2002), and the ***B2M*** immune evasion signature (Zaretsky et al., 2016) — provide highly validated targets for therapeutic disruption. Together, these findings demonstrate the power of integrating explainable artificial intelligence with network biology to discover novel biomarkers and therapeutic targets in precision oncology.

## CODE AVAILABILITY

The modular Python execution workflow implementing the preprocessing, XGBoost classification, SHAP explainability, LIANA inference, and survival analysis algorithms is freely available for academic review and reproduction. The complete 10-cell Google Colab-compatible pipeline will be deposited in a public GitHub repository upon acceptance.

## AUTHOR CONTRIBUTIONS

**R.O**.: Pipeline design, implementation, data analysis, figure generation, manuscript drafting. **E.N**.: Project supervision, conceptualisation, critical revision of manuscript. Both authors have read and approved the final version.

